# TernTables: A Statistical Analysis and Table Generation Web Interface for Clinical and Biomedical Research

**DOI:** 10.64898/2026.04.15.717241

**Authors:** Joshua D. Preston, Helen Abadiotakis, Ailin Tang, Clayton J. Rust, Michael E. Halkos, Mani A. Daneshmand, Joshua L. Chan

**Author notes:** **Corresponding Author:** Joshua L. Chan, MD, Carlyle Fraser Cardiothoracic Surgery Research Laboratory, Division of Cardiothoracic Surgery, Department of Surgery, Emory University School of Medicine, 1365 Clifton Rd NE, Atlanta, GA 30322.

## Abstract

Clinical research dissemination is frequently hindered by administrative friction and methodological inconsistency. To address these barriers, we developed TernTables, a freely available, open-source web application (https://www.tern-tables.com/) and R package (https://cran.r-project.org/package=TernTables) that streamlines the transition from raw data to formatted results for descriptive and univariate clinical reporting. The system integrates a client-side screening protocol for protected health information (PHI) with a rule-based decision tree that selects and executes appropriate frequency-based, parametric, or non-parametric statistical tests based on data distribution and class. TernTables generates publication-ready summary tables in Microsoft Word format, complemented by dynamically generated methods text and the underlying R code to ensure complete transparency and reproducibility. Validation using a landmark clinical trial dataset demonstrated concordance with established biostatistical approaches for descriptive and univariate analyses. TernTables is designed to supplement, not replace, formal statistical consultation by standardizing routine descriptive and univariate workflows, allowing biostatistical expertise to be focused on complex analyses and study design. By lowering technical and financial barriers, the platform democratizes access to rigorous statistical workflows while maintaining methodological excellence and reducing “researcher degrees of freedom.”

## INTRODUCTION

Clinical research dissemination is hindered by two barriers: administrative friction and methodological inconsistency. This discourages surgeons from initiating research and can introduce variability and reduce reproducibility.

Although data synthesis and summary tables are foundational for reporting clinical research, “extra paperwork” is cited as the primary deterrent to clinician-led research by 77% of physicians.^1^ Dedicated statistical expertise shortens publication timelines,^2^ yet competing demands on institutional biostatisticians often prioritize complex modeling over simpler analyses. Documented gaps in physician biostatistical literacy^3^ leave clinicians ill-equipped to perform basic analyses. Most importantly, the absence of standardized workflows introduces “researcher degrees of freedom,” allowing subjective analytic choices to produce divergent results from identical datasets,^4^ a problem not fixed by editorial guidance on statistical reporting alone.^5^ Such variability can alter observed treatment effects by nearly 40% and contribute to misinterpretation.^6^ Importantly, many early-stage or exploratory clinical questions rely primarily on descriptive and univariate analyses, which clinicians may appropriately conduct when supported by standardized, transparent workflows.

To address these challenges, we developed TernTables, a freely available, open-source web application and R package that expedites the clinical research reporting workflow. The system integrates raw-data preprocessing with rule-based statistical testing to generate publication-ready summary tables for descriptive and univariate analyses. Unlike existing statistical software that requires programming expertise or manual formatting, TernTables integrates preprocessing, systematic test selection, and formatted table generation with a no-code interface. The interface executes reproducible R-based computation, bridging the gap between accessibility and computational rigor. By lowering time and expertise barriers, TernTables lets clinicians focus on clinical inquiry while maintaining methodological rigor.

## METHODS

The TernTables web application (https://www.tern-tables.com/) is an accessible interface for its underlying R package engine (https://cran.r-project.org/package=TernTables). The system accepts tabular datasets with patient-level observations. To prioritize data security, a client-side screening protocol blocks uploads containing flagged terms associated with protected health information (PHI).

The core architecture uses a hierarchical decision tree to standardize summary-statistic formatting and select appropriate statistical tests for univariate analysis (**Table 1**). Following preprocessing, variables undergo a multi-gate normality assessment incorporating skewness, kurtosis, sample size, and the Shapiro-Wilk test to route continuous variables to parametric (Welch t-test or Welch ANOVA) or non-parametric (Wilcoxon rank-sum or Kruskal-Wallis) methods; categorical variables are evaluated using Pearson χ^2^ or Fisher exact test based on expected cell counts. The system then generates publication-ready tables in Microsoft Word format, accompanied by dynamically generated methodological text documenting each statistical test applied and a copy of all executed R code.

**Table 1.** Rule-based statistical decision framework for standardized clinical reporting enforced by TernTables.

| Variable Class and Distribution | 2 Groups | 3 or More Groups (Omnibus) | Post-hoc Analysis<sup>a</sup> |
| --- | --- | --- | --- |
| Continuous, Normal<sup>b</sup> | Welch *t*-test | Welch ANOVA | Games-Howell |
| Continuous, Non-normal<sup>b</sup> | Wilcoxon rank sum | Kruskal-Wallis | Dunn test + Holm |
| Ordinal<sup>c</sup> | Wilcoxon rank sum | Kruskal-Wallis | Dunn test + Holm |
| Categorical<sup>d</sup> | $\chi^2$ or Fisher exact | $\chi^2$ or Fisher exact | Adjusted standardized residuals |

Abbreviations: ANOVA, analysis of variance
<sup>a</sup> Post-hoc analyses are performed only when the omnibus $P < .05$ .
<sup>b</sup> A hierarchical decision tree determines distribution. Variables are routed to non-parametric tests if any group $n < 3$ , any group absolute skewness $> 2.0$ , or any group absolute excess kurtosis $> 7$ ; variables are routed to parametric tests if all groups $n \geq 30$ (Central Limit Theorem); otherwise, variables are routed to parametric tests if all groups pass the Shapiro-Wilk test ( $P > .05$ ) and to non-parametric tests if any group fails.
<sup>c</sup> Non-parametric routing is enforced when a variable is designated as ordinal to preserve rank-based relationships.
<sup>d</sup> Fisher exact test is enforced when any expected cell count is $< 5$ (Cochran criteria); otherwise, Pearson $\chi^2$ is utilized. When the exact algorithm cannot complete (e.g., tables larger than $2 \times 2$ ), Fisher exact with Monte Carlo simulation ( $B = 10,000$ ) is applied automatically.

## RESULTS

The application was evaluated using a landmark colon cancer trial dataset.^7^ During upload, the security protocol identified and rejected synthetic PHI fields. When challenged with degraded data, the preprocessing engine standardized missingness indicators (e.g., “unk”, “NA”), removed empty structural elements, corrected inconsistent capitalization, and produced a diagnostic audit.

For descriptive summaries, the system reported mean ± SD or median [IQR] based on the normality decision tree. For two- and three-group comparisons, variables were routed to appropriate frequency-based, parametric, or non-parametric tests. The final output demonstrated results consistent with standard biostatistical approaches. Generated tables included summary statistics, *P* values (unadjusted and false discovery rate [FDR]-corrected), unadjusted odds ratios, and *post hoc* comparisons (**Supplemental Content**). A schematic of the program’s end-to-end pipeline is displayed in **Figure 1**.

**Figure 1.**
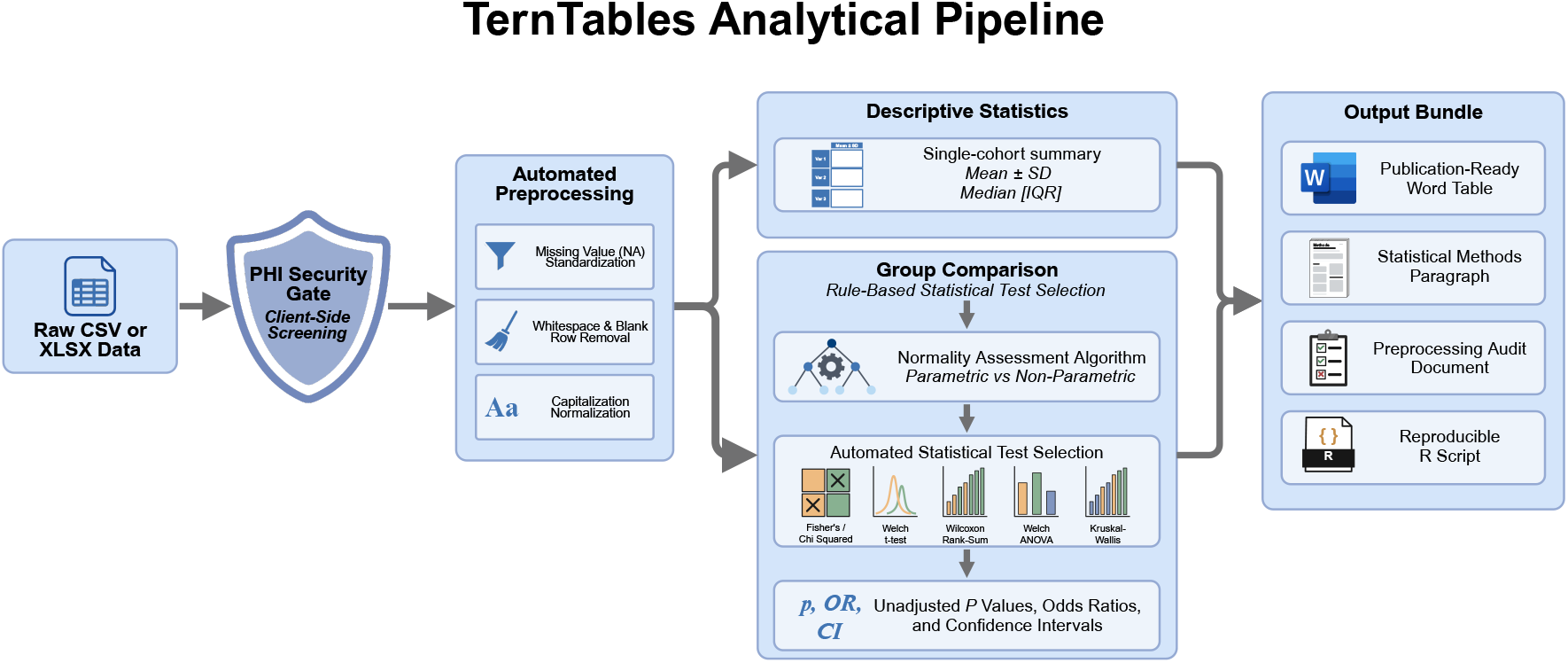
The TernTables end-to-end analytical workflow.

## DISCUSSION

TernTables streamlines the transition from raw data to formatted results, addressing both administrative and methodological barriers that deter physician-led research.

Systematizing the production of publication-ready tables streamlines analytical workflows and provides a standardized framework for routine clinical reporting. This capability is important because limited statistical support bottlenecks surgical research, despite evidence that collaboration with a statistician accelerates publication timelines.^2^ Crucially, TernTables is not intended to offload critical statistical reasoning or replace formal consultation; rather, by enabling investigators to handle routine descriptive and comparative analyses independently, the platform reserves specialized biostatistical collaboration for complex modeling, causal inference, and study design.

In addition to improving efficiency, TernTables provides safeguards against common statistical pitfalls. The platform supports appropriate reporting of baseline characteristics in randomized trials by facilitating descriptive summaries without unnecessary hypothesis testing, thereby addressing the pervasive “Table 1 fallacy.”^8^ For exploratory analyses involving large numbers of variables, the optional FDR adjustment helps mitigate spurious findings. The rule-based decision framework also standardizes statistical test selection, reducing subjective analytic choices that contribute to “researcher degrees of freedom” and inconsistent results.^4,6^ Transparency is further supported through a “glass box” architecture, allowing users to inspect the underlying R code executed during analysis. This design intentionally promotes reproducibility, aligning with principles of responsible conduct of research by ensuring that data analysis tools and code are disclosed with sufficient detail for independent reproduction.^9^

Ultimately, TernTables expands access to standardized statistical workflows through an accessible yet methodologically robust tool that converts raw datasets into reproducible, publication-ready analyses. By lowering analytical barriers and enforcing best practices, the platform encourages broader participation in clinical research and promotes transparency, standardization, and reproducibility.

## Supporting information

Supplemental Content

## ACKNOWLEDGEMENTS

The authors would like to thank Maria V. Aslam, PhD, MPH (Department of Surgery, Emory University School of Medicine), for her thorough review of the statistical methods implemented into TernTables.

JLC was supported in part by the National Center for Advancing Translational Sciences of the National Institutes of Health (NIH) under Award Numbers UL1TR002378 and KL2TR002381, as well as the Carlyle Fraser Heart Center. JDP was supported by the NIH Medical Scientist Training Program under Award Number T32GM008169. The authors have no conflicts of interest to declare.

In preparing this work, the authors utilized Grammarly and Google’s Gemini 3 Pro for general editorial assistance, manuscript structuring, and wording guidance. OpenAI’s ChatGPT-5.2 and Gemini 3 Pro were used to identify relevant literature. Anthropic’s Claude Sonnet 4.6 via GitHub Copilot was used to develop, debug, and revise the source code for the R package and web app. The base schematic summary figure was created using PaperBanana^10^ and further refined using BioRender (see https://biorender.com/hybx65h for publication license). The authors reviewed and edited all content and take full responsibility for the publication.

## DATA SHARING STATEMENT

All source code and example datasets for the TernTables R package are publicly available at https://github.com/jdpreston30/TernTables, https://cran.r-project.org/package=TernTables, and https://cran.r-project.org/package=survival.

