## Supplemental Content for "TernTables: A Statistical Analysis and Table Generation Web Interface for Clinical and Biomedical Research"

***Supplemental Table 1. A descriptive summary table generated by TernTables.***

| Category<br>Variable | Total<br>(N = 929) |
|---|---|
| **Patient Demographics** | |
| Age (yr) | 59.8 ± 11.9 |
| Sex | |
| *Female* | 445 (48%) |
| *Male* | 484 (52%) |
| **Surgical Findings** | |
| Colonic Obstruction | 180 (19%) |
| Bowel Perforation | 27 (3%) |
| **Tumor Characteristics** | |
| Positive Lymph Nodes (n) | 2 [1–5] |
| > 4 Positive Nodes | 255 (27%) |
| Tumor Adherence | 135 (15%) |
| Tumor Differentiation | |
| *Well* | 93 (10%) |
| *Moderate* | 663 (73%) |
| *Poor* | 150 (17%) |
| Extent Of Local Spread | |
| *Submucosa* | 21 (2%) |
| *Muscle* | 106 (11%) |
| *Serosa* | 759 (82%) |
| *Contiguous Structures* | 43 (5%) |
| **Outcomes** | |
| Recurrence | |
| *No Recurrence* | 461 (50%) |
| *Recurrence* | 468 (50%) |
| Treatment Arm | |
| *Levamisole + 5FU* | 304 (33%) |
| *Levamisole* | 310 (33%) |
| *Observation* | 315 (34%) |

***Supplemental Table 2. A two-group comparison table generated by TernTables.***

| Category<br>Variable | No Recurrence (n = 461) | Recurrence (n = 468) | Total (N = 929) | OR | P value |
|---|---|---|---|---|---|
| **Patient Demographics** | | | | | |
| Age (yr) | 60.5 ± 11.5 | 59 ± 12.4 | 59.8 ± 11.9 | - | 0.065 |
| Sex | | | | | |
| *Female* | 216 (47%) | 229 (49%) | 445 (48%) | 1.00 (ref.) | 0.570 |
| *Male* | 245 (53%) | 239 (51%) | 484 (52%) | 0.92 [0.71–1.19] | - |
| **Surgical Findings** | | | | | |
| Colonic Obstruction | 81 (18%) | 99 (21%) | 180 (19%) | 1.26 [0.91–1.75] | 0.194 |
| Bowel Perforation | 10 (2%) | 17 (4%) | 27 (3%) | 1.70 [0.77–3.75] | 0.258 |
| **Tumor Characteristics** | | | | | |
| Positive Lymph Nodes (n) | 2 [1–3] | 3 [2–6] | 2 [1–5] | - | **2E-14** |
| > 4 Positive Nodes | 75 (16%) | 180 (38%) | 255 (27%) | **3.22 [2.36–4.38]** | **6E-14** |
| Tumor Adherence | 53 (11%) | 82 (18%) | 135 (15%) | **1.64 [1.13–2.37]** | **0.012** |
| Tumor Differentiation | | | | | |
| *Well* | 49 (11%) | 44 (10%) | 93 (10%) | - | 0.089 |
| *Moderate* | 337 (75%) | 326 (71%) | 663 (73%) | - | - |
| *Poor* | 62 (14%) | 88 (19%) | 150 (17%) | - | - |
| Extent Of Local Spread | | | | | |
| *Submucosa* | 16 (3%) | 5 (1%) | 21 (2%) | - | **7E-6** |
| *Muscle* | 72 (16%) | 34 (7%) | 106 (11%) | - | - |
| *Serosa* | 359 (78%) | 400 (85%) | 759 (82%) | - | - |
| *Contiguous Structures* | 14 (3%) | 29 (6%) | 43 (5%) | - | - |
| **Treatment Details** | | | | | |
| Treatment Arm | | | | | |
| *Levamisole + 5FU* | 185 (40%) | 119 (25%) | 304 (33%) | - | **1E-5** |
| *Levamisole* | 138 (30%) | 172 (37%) | 310 (33%) | - | - |
| *Observation* | 138 (30%) | 177 (38%) | 315 (34%) | - | - |

***Supplemental Table 3. A three-group comparison table generated by TernTables.***

| **Category**<br>**Variable** | **Observation (n = 315)** | **Levamisole (n = 310)** | **Levamisole + 5FU (n = 304)** | **Total (N = 929)** | **P value** |
|---|---|---|---|---|---|
| **Patient Demographics** | | | | | |
| Age (yr) | 59.5 ± 12 | 60.1 ± 11.6 | 59.7 ± 12.3 | 59.8 ± 11.9 | 0.780 |
| Sex | | | | | |
| *Female* | 149 (47%) | 133 (43%) | 163 (54%) | 445 (48%) | **0.028** |
| *Male* | 166 (53%) | 177 (57%) | 141 (46%) | 484 (52%) | - |
| **Surgical Findings** | | | | | |
| Colonic Obstruction | 63 (20%) | 63 (20%) | 54 (18%) | 180 (19%) | 0.683 |
| Bowel Perforation | 9 (3%) | 10 (3%) | 8 (3%) | 27 (3%) | 0.907 |
| **Tumor Characteristics** | | | | | |
| Positive Lymph Nodes (n) | 2 [1–5] | 2 [1–5] | 2 [1–4] | 2 [1–5] | 0.550 |
| > 4 Positive Nodes | 87 (28%) | 89 (29%) | 79 (26%) | 255 (27%) | 0.749 |
| Tumor Adherence | 47 (15%) | 49 (16%) | 39 (13%) | 135 (15%) | 0.562 |
| Tumor Differentiation | | | | | |
| *Well* | 27 (9%) | 37 (12%) | 29 (10%) | 93 (10%) | 0.522 |
| *Moderate* | 229 (74%) | 219 (73%) | 215 (72%) | 663 (73%) | - |
| *Poor* | 52 (17%) | 44 (15%) | 54 (18%) | 150 (17%) | - |
| Extent Of Local Spread | | | | | |
| *Submucosa* | 8 (3%) | 3 (1%) | 10 (3%) | 21 (2%) | 0.264 |
| *Muscle* | 38 (12%) | 36 (12%) | 32 (11%) | 106 (11%) | - |
| *Serosa* | 249 (79%) | 259 (84%) | 251 (83%) | 759 (82%) | - |
| *Contiguous Structures* | 20 (6%) | 12 (4%) | 11 (4%) | 43 (5%) | - |
| **Outcomes** | | | | | |
| Recurrence | | | | | |
| *No Recurrence* | 138 (44%) | 138 (45%) | 185 (61%) | 461 (50%) | **1E-5** |
| *Recurrence* | 177 (56%) | 172 (55%) | 119 (39%) | 468 (50%) | - |
